# Which lower-limb joints compensate for destabilising energy during walking in humans?

**DOI:** 10.1101/2022.01.11.475955

**Authors:** Pawel R. Golyski, Gregory S. Sawicki

## Abstract

Maintaining stability during perturbed locomotion requires coordinated responses across multiple levels of organization (e.g., legs, joints, muscle-tendon units). However, current approaches to investigating such responses lack a “common currency” that is both shared across scales and can be directly related to perturbation demands. We used mechanical energetics to investigate the demands imposed on a leg by a transient increase in unilateral treadmill belt speed targeted to either early or late stance. We collected full body kinematics and kinetics from 7 healthy participants during 222 total perturbations. From across-subject means, we found early stance perturbations elicited no change in net work exchanged between the perturbed leg and the treadmill but net positive work at the overall leg level, and late stance perturbations elicited positive work at the leg/treadmill interface but no change in net work at the overall leg level. Across all perturbations, changes in ankle and knee work from steady state best reflected changes in overall leg work on the perturbed and contralateral sides, respectively. Broadening this paradigm to include joint level (vs. leg level) perturbations and including muscle-tendon unit mechanical energetics may reveal neuromechanical responses used in destabilizing environments which could inform design of balance-assisting devices and interventions.

**Subject Areas:** biomechanics, biomedical engineering, bioengineering

## 1. Introduction

Falls remain a major public health problem. In the United States alone, 1 in 4 adults over 65 years old fall at least once a year, which result in over 25,000 deaths annually and $31 billion in annual direct healthcare costs [1–3]. In the workplace, falls caused 16% of fatal work-related injuries in 2019, with 68% of fall-related injuries occurring in individuals between 35 and 64 years old [4,5]. In both younger and older adults, falls occur more often during walking than any other locomotor task, with external disturbances such as slips and trips being the predominant perceived cause of falls [6,7]. Although responses to external perturbations have been extensively studied at the scales of the overall leg (i.e., foot placement), joints, and muscles (e.g., [8–17]), how stabilizing responses are related and coordinated across scales is not well understood. Two obstacles to such analyses are: 1) variables used to characterise responses are generally not measured using a “common currency” that can be easily related across different scales, and 2) the explicit, quantifiable demand imposed by the perturbation is unknown. In this work we aimed to overcome these obstacles by using a split-belt treadmill to deliver destabilizing perturbations using transient changes in belt speed that imposed quantifiable energetic demands on the legs that could be related to changes in work at the joints. We anticipate this analysis will serve as an initial step in multi-scale analysis of stability through mechanical energetics.

The mechanical power of each leg during walking, with respect to a fixed global reference frame and assuming massless legs, can be estimated using the individual limbs method [18], which quantifies the mechanical energy flowing between the ground and the centre of mass (COM). During overground walking, the mechanical power of each leg is the dot product of its corresponding ground reaction force (GRF) and COM velocity. No power flows between each leg and the ground because the velocity of the ground is 0, thus ***F***_*Leg*_ · ***v***_*ground*_ =, where ***F***_*Leg*_ is equal and opposite to the ground reaction force. However, for treadmill walking, this is no longer the case; with respect to a fixed global reference frame, each belt is moving, so the power flowing from each leg to its corresponding belt is ***F***_*Leg*_ · ***v***_*belt*_. Thus, in the case of treadmill walking, the mechanical power of each leg is the sum of the power flowing from the leg to the treadmill belt and the leg to the COM [19].

During level ground treadmill walking with both belts of a split-belt treadmill moving at the same speed, the net work of each leg on the COM is zero on average over a stride. Further, with both belts moving at the same speed, since the average anteroposterior force must be zero over a stride (otherwise the COM would accelerate relative to the treadmill), each leg performs zero net mechanical work on its corresponding belt. Therefore, in this condition the net work performed by each leg must be zero on average [20]. However, in either non-steady conditions or when belts of a split-belt treadmill are moving at different speeds, the previous assumptions no longer hold, and a treadmill can elicit an energetic demand on the leg over a stride. In this work, we leveraged this concept and designed perturbations intended to elicit a change in net work over a stride by a leg (i.e. generation or dissipation). Specifically, since leg force is directed anteriorly in early stance and posteriorly in late stance, by increasing the posterior velocity of a belt our first hypothesis was net negative work would be elicited over a stride at the leg/treadmill interface with an early stance perturbation, while net positive work would be elicited over a stride with a late stance perturbation (figure 1). Our second hypothesis was that such changes in net work at the leg/treadmill interface relative to an unperturbed stride would be reflected at the level of overall leg work, since the treadmill environment limits large fluctuations in COM velocity and thus mechanical power exchanged between each leg and the COM (***F***_*GRF*_ · ***v***_*COM*_).

**Figure 1.**
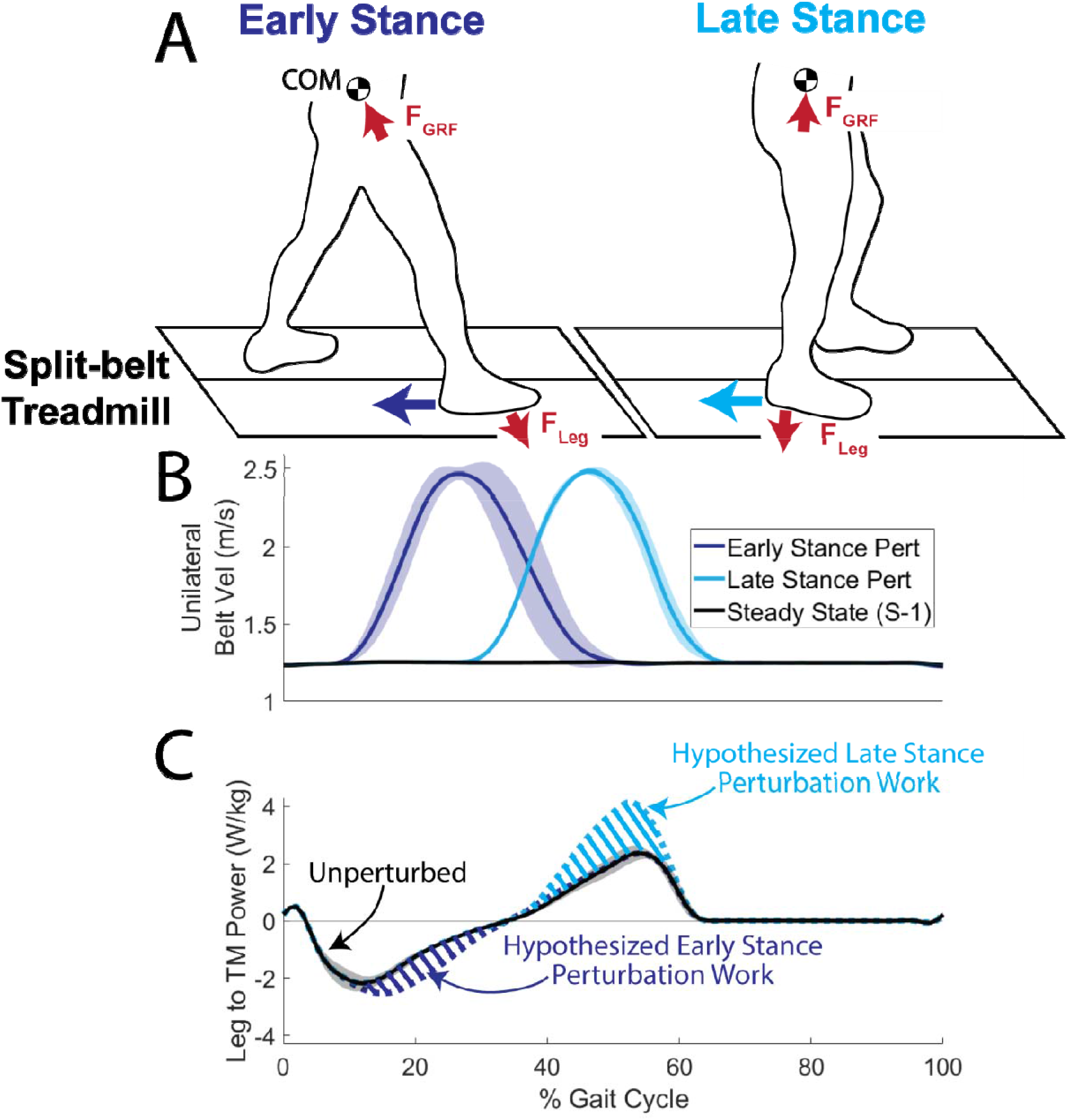
Perturbation overview. A) Perturbation design. In early stance perturbations, since the leg force is anterior and treadmill velocity is posterior, this results in the leg performing negative work on the belt. In late stance, when leg force is posterior and treadmill velocity is posterior, the leg performs positive work on the belt. B) Unilateral belt velocity traces show across-subject ensemble averages. Shaded regions represent ±1 standard deviation. C) Hypothesized changes in power flowing from the leg to the treadmill relative to steady state, with early stance perturbations eliciting negative power (e.g., dissipation) and late stance perturbations eliciting positive power (e.g., generation).

While no previous work has investigated the mechanical energetics of transient unilateral treadmill speed perturbations during walking, there are numerous experimental contexts that elicit an energetic demand on the legs during locomotion by changing the required amount of work the legs perform on the COM. Examples include increasing or decreasing the slope of the ground relative to level [21–25], accelerating or decelerating [26,27], and falling into a hole [28,29]. In general, relative to level ground, walking on an incline results in a shift to more positive work and power at the hip joint [21–24], while walking on a decline results in more negative work and power at the knee joint [22–24]. However, in the case of speed changes and ground height perturbations, changes in ankle work and power best reflect the overall demand on the leg [26–30]. Since our perturbations were more similar to speed changes than slope changes, our third hypothesis was that changes in net leg work over a stride resulting from the perturbation would primarily be reflected by changes in net work at the ankle joint.

## 2. Methods

### 2.1 Experimental protocol

Seven young, healthy individuals (5 males, 2 females, mean [SD]: 25 [2] years, 178.5 [12.1] cm stature, 72.7 [13.3] kg) walked on an instrumented split-belt treadmill (CAREN; Motek, Netherlands). Following an unperturbed 5-minute acclimation period at 1.25 m/s [31], transient unilateral belt accelerations were delivered during walking. Each perturbation was targeted to either early or late stance and either the left or right leg. Each timing/leg pairing was repeated 10 times (i.e., 2 legs x 2 timings x 10 repetitions = 40 perturbations per participant). The order of perturbations was randomized, with 30-40 steps between perturbations to ensure the perturbation was unexpected and the participant had returned to steady state walking [32]. The perturbation algorithm is fully described elsewhere [33], and used real-time kinematic data to estimate gait phase during walking. Perturbations consisted of a brief (mean duration: 340 ms, 32.9% perturbed gait cycle) increase in belt speed from 1.25 m/s to 2.5 m/s (figure 1B). All participants provided informed consent as approved by the local Institutional Review Board.

### 2.2 Kinematics and kinetics

67 reflective markers (modified Human Body Model 2; [34]) were placed on the bony landmarks and major body segments (head, hands, forearms, upper arms, torso, pelvis, thighs, shanks, and feet) of each participant. A 10-camera motion capture system (Vicon; Oxford, UK) collected 3D marker trajectories at 200 Hz. For each participant, a static trial was used to scale an individualized version of the generic full body musculoskeletal model developed by Rajagopal and colleagues (37 degrees of freedom, 22 rigid bodies; [35]). The metatarsophalangeal and subtalar joints of the models were locked in all analyses, thereby assuming the foot was a rigid body. These subject specific models, in conjunction with measured marker positions, were used to calculate joint angles for each trial using the inverse kinematics tool in OpenSim v. 4.0 [36]. Joint moments were calculated with the inverse dynamics tool in OpenSim using both the joint angles and bilateral ground reaction forces (2000 Hz sampling) applied to the calcanei of the scaled models. Joint kinematics and kinetics were lowpass filtered using 4^th^ order zero-phase Butterworth filters at 6 and 15 Hz, respectively. Strides were segmented using a 30 N threshold from vertical ground reaction forces. Trials where participants crossed over the belts, as determined by manual inspection, were removed from analysis, leaving 222 successful trials.

### 2.3 Mechanical energetics

Joint powers were calculated as the product of joint moments and joint angular velocities. To estimate leg power from summed joint power, joint powers about each available degree of freedom were summed (ankle plantar/dorsiflexion, knee flexion/extension, hip flexion/extension, hip ab/adduction, and hip external/internal rotation). To estimate leg power most accurately from joint power, 6 degree of freedom joint powers and a deformable segment model for the foot are preferred [37–40], but such calculations are not compatible with model-based OpenSim analysis that constrains joints to behave within physiological bounds. Thus, an additional “ground-truth” estimate of leg power was calculated using a modified individual limbs method [18,19,37]. This corrected leg power was the sum of 1) the power flowing from each leg to the treadmill, 2) the power flowing from each leg to the COM, and 3) the peripheral power of the leg segments (thigh, shank, and foot) relative to the COM. The power from the leg to the treadmill was calculated as the dot product of the force from each leg on the ground (equal and opposite to ground reaction force) with the velocity of the respective treadmill belt (logged at approximately 70 Hz). The power from each leg to the COM was calculated as the dot product of each ground reaction force and the velocity of the whole-body COM as calculated by the Body Kinematics tool in OpenSim. This kinematic COM estimate was selected instead of estimates based on GRF [18–20] because GRF-based estimates of COM velocity solve for integration constants assuming steady-state behaviour over multiple strides, but such integration constants may not be valid during perturbed strides. Peripheral power of the leg segments was calculated by summing the time derivative of the rotational and translational components of kinetic energy [37,41,42]. Mechanical work at the levels of the joints and legs was calculated using trapezoidal integration of joint powers.

### 2.4 Statistical analysis

All statistical analyses were performed in Matlab R2019b (Mathworks, Natick, MA, USA). To characterize the influence of the perturbation, sagittal plane extrema in kinematic and kinetics, in addition to mechanical work estimates, were evaluated using a linear mixed model with fixed effects of perturbation timing (2 levels: early and late) and ipsilateral stride in relation to the perturbation [4 levels: S-1 (the preceding unperturbed, steady state stride), S0 (perturbed stride), S+1 (first recovery stride), S+2 (second recovery stride)], in addition to a random effect of participant. Bonferroni-corrected pairwise t-tests were run for post-hoc analyses. Linear regressions were used to relate changes in corrected leg work with changes in ankle, knee, and hip work. For these regressions, changes in leg and joint work were calculated relative to steady state strides (i.e., the work of stride S+N – work of stride S-1). Significance was concluded for p-values ≤ 0.05.

## 3. Results

### 3.1 Peak joint angles

Significant timing x stride interactions were observed in ipsilateral ankle, knee, and hip angles, in addition to contralateral ankle and knee angles (electronic supplementary material, table S1). Differences in kinematics following the perturbation relative to steady state (S-1) were predominantly limited to the perturbed stride (S0) and first recovery stride (S+1; figure 2).

**Figure 2.**
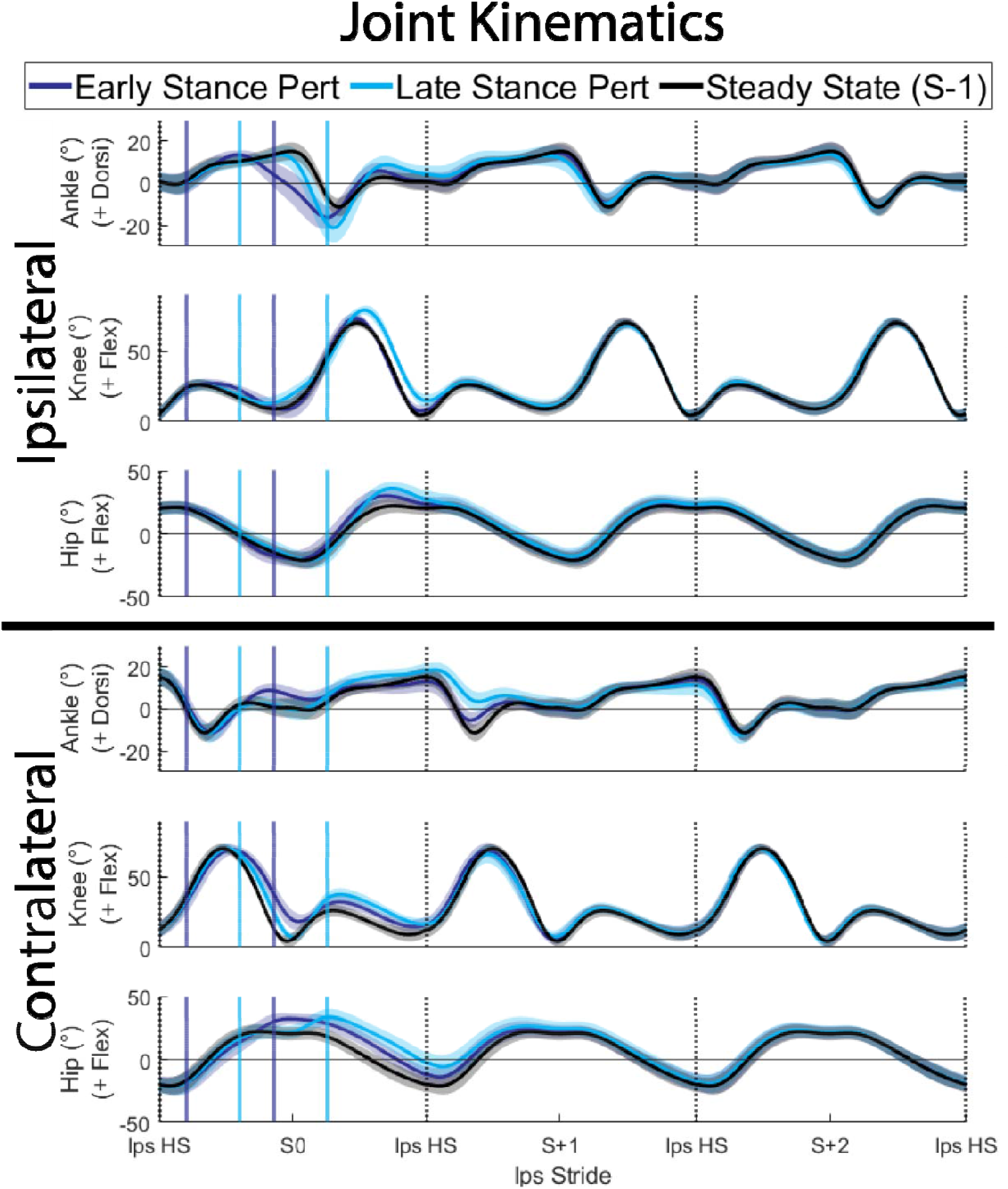
Lower limb joint angles averaged across subjects and normalized to percentage of the gait cycle. Shaded areas represent ±1 standard deviation. Solid vertical lines indicate the average start and end times of the perturbations. “Steady State” strides are the stride preceding the perturbed stride (S-1).

On the perturbed stride (S0) and ipsilateral leg, both early and late stance perturbations resulted in increased plantarflexion (both timings: p < 0.001), knee flexion (early: p = 0.030, late: p < 0.001), and hip flexion (both timings: p < 0.001), with late stance perturbations causing larger deviations than early stance perturbations (p < 0.001 for all). During the perturbed stride (S0) on the contralateral leg there were similar increases in hip flexion for both timings (both timings: p < 0.001). However, early stance perturbations resulted in increased knee flexion at initial contact compared to late stance perturbations, evidenced by a decreased range of motion (p < 0.001), while late stance perturbations resulted in increased dorsiflexion compared to early stance perturbations (p < 0.001).

On the first recovery stride (S+1), increased ipsilateral hip flexion relative to steady state persisted for both timings (early: p = 0.032, late: p < 0.001), with deviations being larger for late vs. early stance perturbations (p < 0.001). Late stance perturbations also resulted in decreased hip extension (p = 0.003). On the contralateral leg, both perturbation timings resulted in decreased plantarflexion (early: p = 0.003, late: p < 0.001) and hip extension (both timings: p < 0.001), with larger deviations in plantarflexion following late compared to early stance perturbations (p < 0.001). Late stance perturbations also resulted in in increased dorsiflexion, decreased knee flexion, and increased hip flexion relative to steady state (p < 0.001 for all).

### 3.2 Peak joint moments

Significant timing x stride interactions were observed in ipsilateral ankle, knee, and hip moments, in addition to contralateral knee extension moments (electronic supplementary material, table S1). Differences in joint moments following the perturbation relative to steady state (S-1) were predominantly limited to the perturbed stride (S0; figure 3).

**Figure 3.**
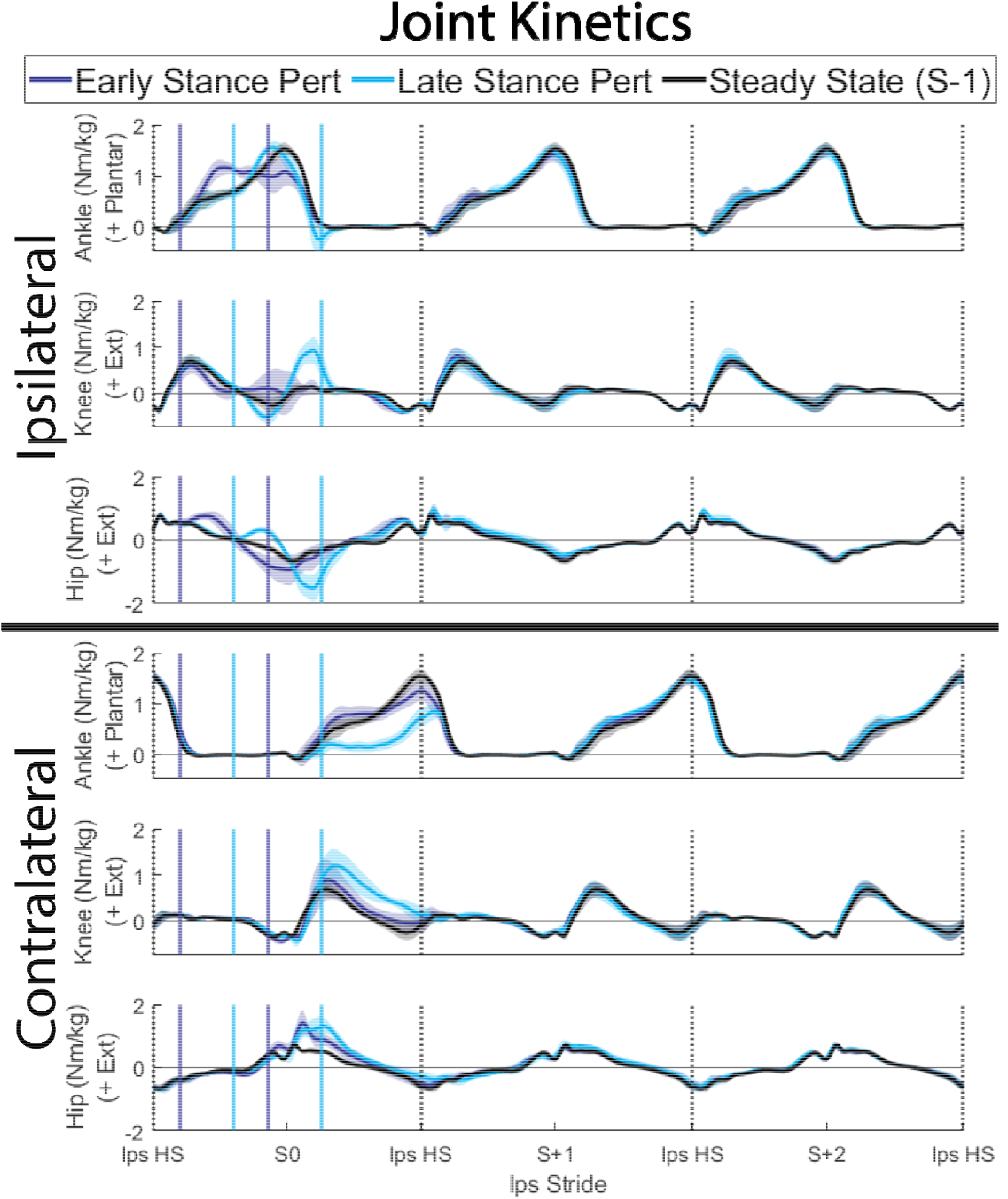
Lower limb joint moments averaged across subjects and normalized to percentage of the gait cycle. Shaded areas represent ±1 standard deviation. Solid vertical lines indicate the average start and end times of the perturbations. “Steady State” strides are the stride preceding the perturbed stride (S-1).

On the perturbed stride (S0) and ipsilateral leg, both early and late stance perturbations resulted in increased knee flexion moments (early: p = 0.013, late: p < 0.001) and hip flexion moments (both timings: p < 0.001), with late stance perturbations resulting in larger flexion moments than early stance perturbations (both joints: p < 0.001). Late stance perturbations also resulted in increased ankle plantarflexion moments (p = 0.002) and knee extension moments (p < 0.001). On the contralateral leg, both early and late stance perturbations resulted in increased knee extension (both timings: p < 0.001), knee flexion (early: p = 0.005, late: p < 0.001), and hip extension moments (both timings: p < 0.001), with late vs. early stance perturbations eliciting larger deviations in knee extension moments (p < 0.001) and knee flexion moments (p = 0.002) from steady state (S-1). Late stance perturbations were also associated with decreased contralateral plantarflexion moments relative to steady state (S-1; p = 0.019).

### 3.3 Peak joint powers

Significant timing x stride interactions were observed in peak mechanical power generation and dissipation at all joints, apart from dissipation at the hip joint (electronic supplementary material, table S1). Similar to joint moments, differences in joint powers following the perturbation relative to steady state (S-1) were predominantly limited to the perturbed stride (S0; figure 4), with the exception of altered power generation at the contralateral ankle in the first recovery stride (S+1).

**Figure 4.**
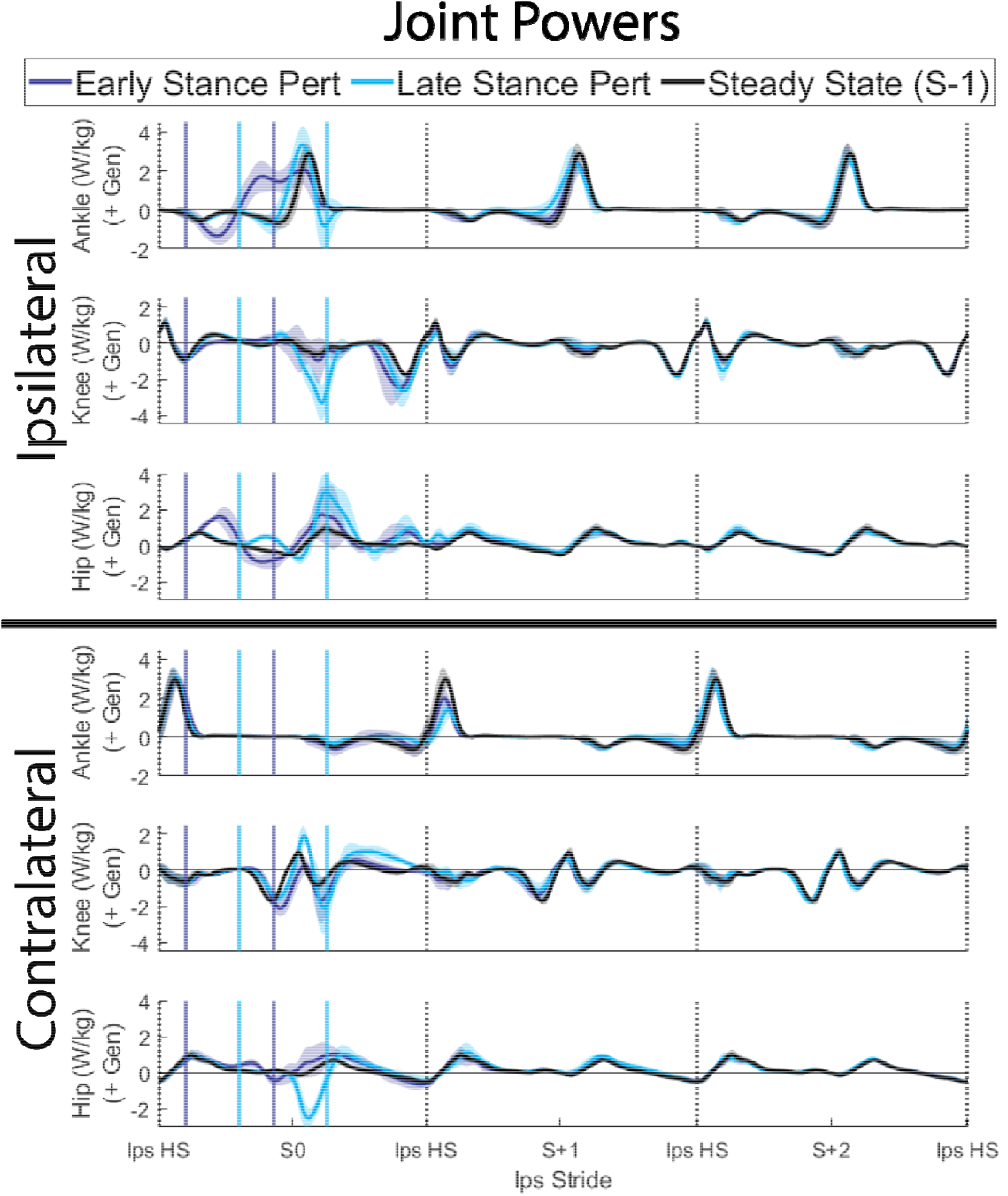
Lower limb joint mechanical powers averaged across subjects and normalized to percentage of the gait cycle. Shaded areas represent ±1 standard deviation. Solid vertical lines indicate the average start and end times of the perturbations. “Steady State” strides are the stride preceding the perturbed stride (S-1).

On the perturbed stride (S0) and ipsilateral leg, both early and late stance perturbations resulted in increased dissipation at the knee (both timings: p < 0.001) and both increased generation and dissipation at the hip (p < 0.001 for all), with late vs. early stance perturbations being associated with increased dissipation at the knee and increased generation at the hip (both joints: p < 0.001). Late stance perturbations additionally resulted in both increased generation and dissipation at the ankle joint (both signs: p < 0.001). The effects of both timings on the ipsilateral hip and knee were similar to the effects on the contralateral joints: both early and late stance perturbations resulted in increased dissipation at the knee (early: p = 0.002, late: p < 0.001), increased dissipation at the hip (both timings: p < 0.001), and increased generation at the hip (early: p < 0.001, late: p = 0.015), with there being more dissipation at the hip in late vs. early stance perturbations (p < 0.001). Further, early stance perturbations resulted in increased dissipation at the contralateral ankle, while late stance perturbations elicited increased generation at the contralateral knee (both joints: p < 0.001).

On the first recovery stride (S+1), the primary differences from steady state (S-1) were decreased generation at the contralateral ankle (both timings: p < 0.001), with less generation for late vs. early stance timings (p = 0.002), in addition to decreased generation at the ipsilateral knee for late stance perturbations (p = 0.011).

### 3.4 Leg mechanical energetics

On the perturbed stride (S0) and ipsilateral leg, corrected leg powers better matched summed joint powers for early vs. late stance perturbations (figure 5A, R^2^ = 0.82 for early, R^2^ = 0.54 for late). Relative to steady state walking (S-1), early stance perturbations elicited more corrected leg work from the perturbed leg on the perturbed stride (p = 0.001), while late stance perturbations did not elicit a significant change in corrected leg work (p = 1.000; figure 5C). For early stance perturbations, this overall increase in corrected leg work was mediated by an increase in work performed by the ipsilateral leg on the COM (p < 0.001), while work performed by the leg on the treadmill was unchanged (p = 1.000). For late stance perturbations, the unchanged overall work performed by the ipsilateral leg was due to increased negative work performed by the leg on the COM (p < 0.001) being cancelled by more positive work being performed by the leg on treadmill (p = 0.005). There was no change in peripheral leg work for either perturbation timing (p > 0.405).

**Figure 5.**
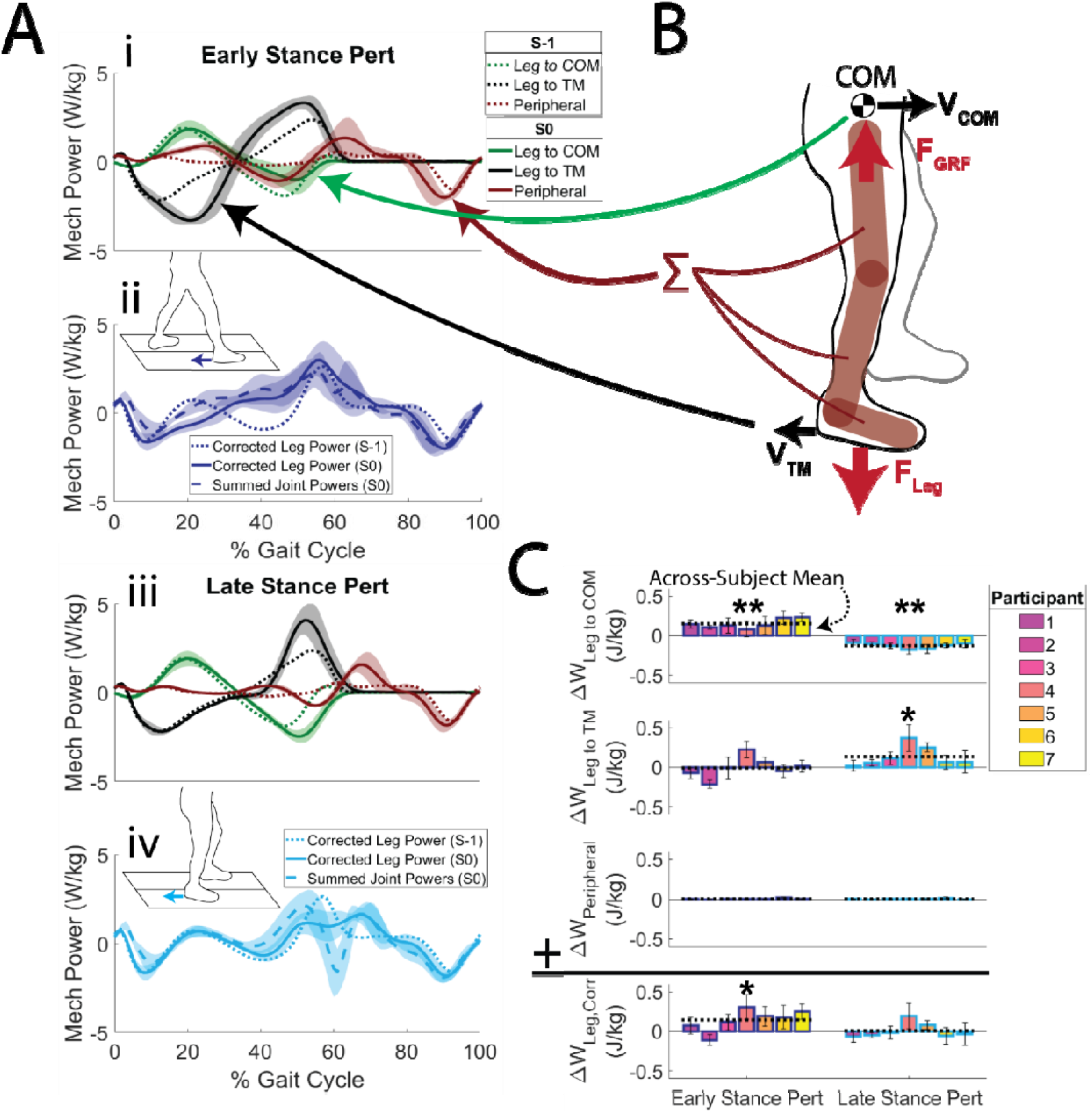
A) Leg mechanical power breakdown for early stance (i and ii) and late stance (iii and iv) perturbations on the ipsilateral leg during the perturbed stride (S0, solid lines), and the unperturbed stride (S-1, dotted lines). For each perturbation timing, across-subject ensemble averages of the contributions to corrected individual leg power are shown (i.e., mechanical power flowing from the leg to the centre of mass, power flowing from the leg to the treadmill, and peripheral power of leg segments moving relative to the centre of mass). These summed contributions (i.e., Corrected Leg Power) are compared with estimated leg power from summing available joint powers (ii and iv). Shaded regions represent ±1 standard deviation. B) Graphical representation of contributions to corrected leg power. As described in section 2.3, power flowing for the leg to the centre of mass or treadmill were the dot product of ground reaction force and centre of mass velocity or leg force and treadmill belt velocity, respectively. Peripheral power contributions were calculated by differentiating the energy of leg segments in time. C) Contributions to corrected leg power were integrated in time to calculate the work contributions over the perturbed stride (S0). These values were subtracted from their respective measurements during steady state strides (S-1) to quantify the changes in mechanical work of these contributions because of the perturbation. Error bars represent ±1 standard deviation. Statistical outcomes represent results of pairwise comparisons of across-subject mean work on the S0 vs. S-1 strides within each timing. * = p < 0.05, ** = p < 0.001

Relating individual joint work with corrected leg work suggests that on the perturbed stride, changes in ipsilateral ankle (R^2^ = 0.56, figure 6i) and contralateral knee work (R^2^ = 0.67, figure 6ii) best reflect changes in overall leg work as a result of the perturbation. On the first recovery stride, changes in ipsilateral ankle (R^2^ = 0.60, figure 6iii) and hip work (R^2^ = 0.39, figure 6iv) best reflected changes in corrected leg work.

**Figure 6.**
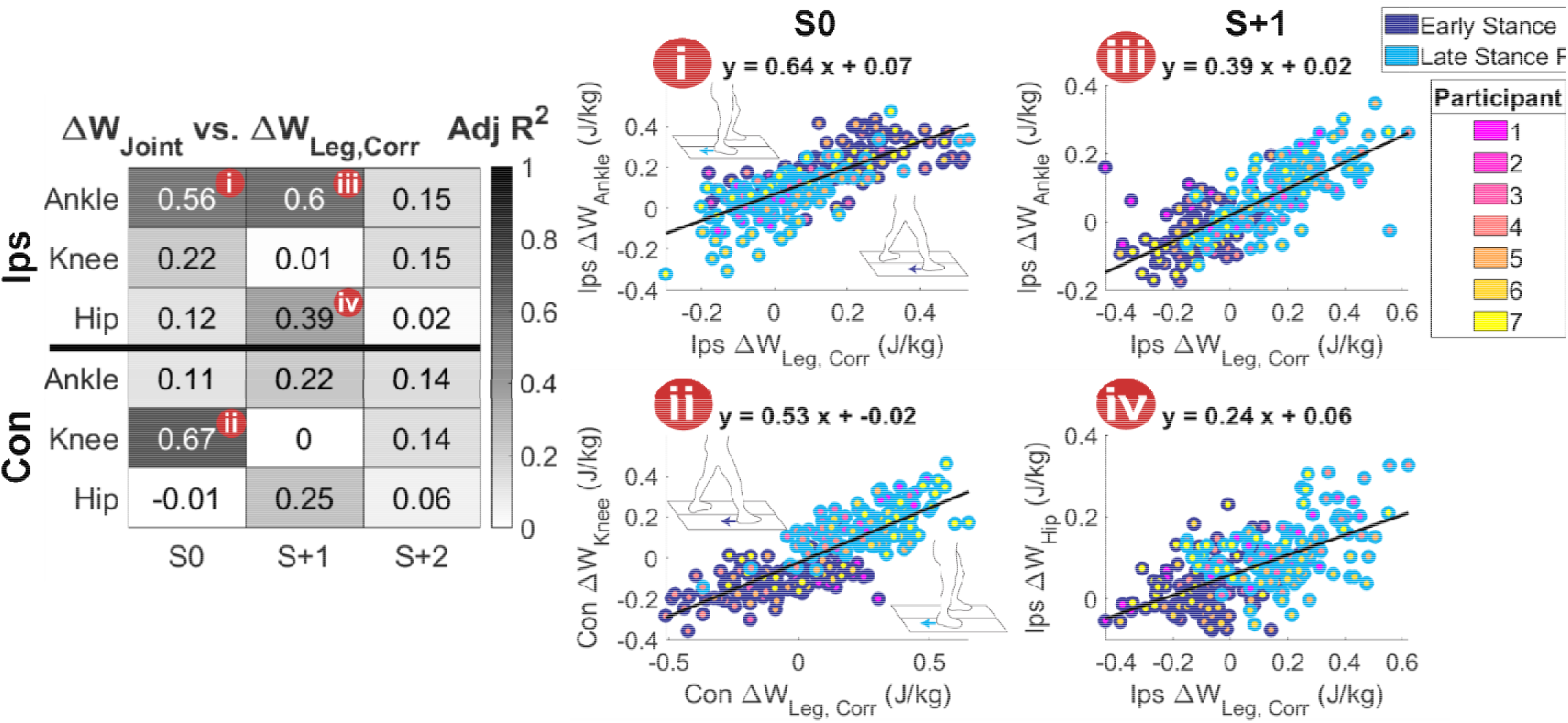
Adjusted R squared values for linear regressions between differences in joint work over a stride from steady state, and differences in corrected leg work over a stride from steady state (S-1) across all 222 perturbations. (i-iv) Scatter plots for joint/stride with the highest adjusted R squared values.

## 4. Discussion

The overall objective of this work was to relate overall leg and joint level responses to destabilizing perturbations during walking using mechanical energetics. We used a split-belt treadmill to elicit transient mechanical energetic demands on the legs during walking and investigated which joints best reflected those demands. Our first hypothesis was that unilateral belt accelerations delivered in early or late stance would elicit net negative or positive work, respectively, from the perturbed leg at the leg/treadmill interface over a stride. Our data supported this hypothesis for late stance perturbations, but not for early stance perturbations. In the case of early stance perturbations, while more negative power was elicited from the leg at the leg/treadmill interface in early stance, the posterior movement of the leg caused by the perturbation led to a more posteriorly directed leg force ([10], electronic supplementary material, figure S1). This posterior leg force resulted in more positive power flowing from the leg to the treadmill in late stance, and no change in net work over a stride by the perturbed leg on the treadmill belt. Thus, future work seeking to specifically elicit net negative work at the leg/treadmill interface over a stride should consider decelerating the targeted treadmill belt during late stance, thereby avoiding unexpected compensations to the perturbation.

Our second hypothesis was that changes in net work at the leg/treadmill interface over the perturbed stride would be reflected by changes in overall leg work. Our data did not support this hypothesis for either early or late stance perturbations. For early stance perturbations, which we initially hypothesized would elicit net negative work from the perturbed leg, we found net positive work was generated by the leg. This occurred due to the combined effect of the net zero work performed by the leg on the treadmill coupled with less negative COM work in late stance. We attribute this decrease in negative work to a combination of 1) the perturbed leg accelerating the COM forward, as evidenced by an increased anteroposterior component of leg power, and 2) offloading of the perturbed leg around toe-off resulting in decreased negative COM power (electronic supplementary material, figure S2). For late stance perturbations, which we initially hypothesized would elicit net positive work from the perturbed leg, we found no net change in work performed by the leg. In this case, the increased positive work performed by the leg on the treadmill was offset by the increased negative work of the leg on the COM in late stance. Increased negative COM power occurred despite the perturbed leg being offloaded, indicating the COM experienced a larger downward velocity around toe-off during late stance perturbations (electronic supplementary material, figure S1 and S2). Since the COM during double support is closer to the ground with faster walking speeds [43], this downward velocity may stem from the increased COM velocity caused by the perturbation coinciding with late stance. An additional observation from the perturbed overall leg mechanical energetic responses was differences among participants, which was driven primarily by differences in work at the leg/treadmill interface (figure 5C). This emphasizes the need for subject specificity in devices or interventions designed to improve perturbation response.

Our third hypothesis was that changes in net ankle work elicited by perturbations would best reflect changes in net overall leg work. Although the energetic demands imposed on the perturbed leg were not as hypothesized, our perturbations nevertheless elicited both generation and dissipation, providing a rich data set to relate leg and joint level mechanical energetics. Our hypothesized contribution by the ankle was supported on the perturbed stride and first recovery stride on the perturbed leg, with a large percentage of the change in work of the perturbed leg being accounted for by the ankle alone (64 and 39% for the perturbed and first recovery stride, respectively). While the importance of the ankle joint in generating mechanical power during steady state walking [21,22,44] and acceleration [26,27,30] has been established, our findings demonstrate that the ankle also plays an important role in mediating transient demands, in agreement with previous studies of human hopping [28]. However, in contrast to the perturbed leg, for the contralateral leg, the knee joint best reflected changes in leg work during the perturbed stride. While previous studies have identified the knee joint as a major contributor during tasks requiring dissipation, such as deceleration and drop landings [27,45], we found that the knee additionally reflected the mechanical work of the leg when generation was required. This could be explained by the knee being a major source of collisional and rebound work in early/midstance, as opposed to the ankle, which primarily contributes later in stance through push-off [37]. Further, previous work that disrupted ankle push-off found that both positive and negative knee energetics were significantly altered [46].

One technical limitation of this work was that joint power contributions did not fully account for leg power, particularly during late stance perturbations. These discrepancies may stem from a combination of 1) the use of musculoskeletal model-based inverse dynamics, as opposed to 6 degree of freedom-based inverse dynamics [37–39], and 2) the foot playing a larger role in late stance perturbations [47]. The latter point could be investigated using a musculoskeletal model with free MTP and subtalar joints or using a deformable segment model [40]. Another important limitation of this work was the use of correlations to relate joint and limb level responses. While this approach suggests which joints reflect demands at the leg level, it does not establish whether those joint level responses cause changes at the leg level. Future work may further investigate the energetic link between joint and leg level responses by perturbing joint energetics and observing leg level responses, perhaps using wearable robots that inject/extract mechanical energy [48–50]. Further, future studies may also use other flavours of perturbations, such as belt decelerations [13,16,51], external pushes [52], or obstacles [8], to determine whether these findings generalize to other unstable contexts. Lastly, inverse dynamics can only quantify net joint powers and does not capture the contributions of muscle-tendon units to energy exchanges across either side of a joint (e.g. coactivation [53]), and between joints (e.g. biarticular muscle tendon units [54]) which could be explored using musculoskeletal simulations, electromyography coupled with in-vivo imaging approaches, and animal models [55–57].

In conclusion, we have demonstrated that a framework using mechanical energetics can be used to investigate joint level contributions to the energetic demand imposed by a transient treadmill-based perturbation during human walking. We found that the net energetic demand on the perturbed leg during the perturbed stride varied depending on the timing of the perturbation, with changes in net leg work stemming from both changes in power flowing from the leg to the COM and from the leg to the treadmill. The varied energetic demands imposed across timings revealed that the ankle best reflected changes in energetics of the perturbed leg on the perturbed and first recovery strides, while the contralateral knee best reflected changes in energetics of the contralateral leg during the perturbed stride. We anticipate this work will serve as an initial step in understanding the multi-scale contributions to whole body behaviour in unstable contexts.

## Ethics

All participants provided written informed consent and all protocols were approved by the Institutional Review Board at the Georgia Institute of Technology (Protocol H20163).

## Data accessibility

Biomechanical data for all participants (N=7) is available at https://sites.gatech.edu/hpl/archival-data-from-publications/

## Authors’ contributions

P.R.G. and G.S.S. conceived of the study and designed the experimental protocol; P.R.G. carried out the experiments, analysed the data, and drafted the manuscript; P.R.G. and G.S.S. edited the manuscript. Both authors gave final approval for publication.

## Acknowledgements

The authors thank Jennifer Leestma for her development of the perturbation program and insightful discussions, in addition to Patrick Kim and Nicholas Swaich for assistance with data collection.

---

**Table S1.**
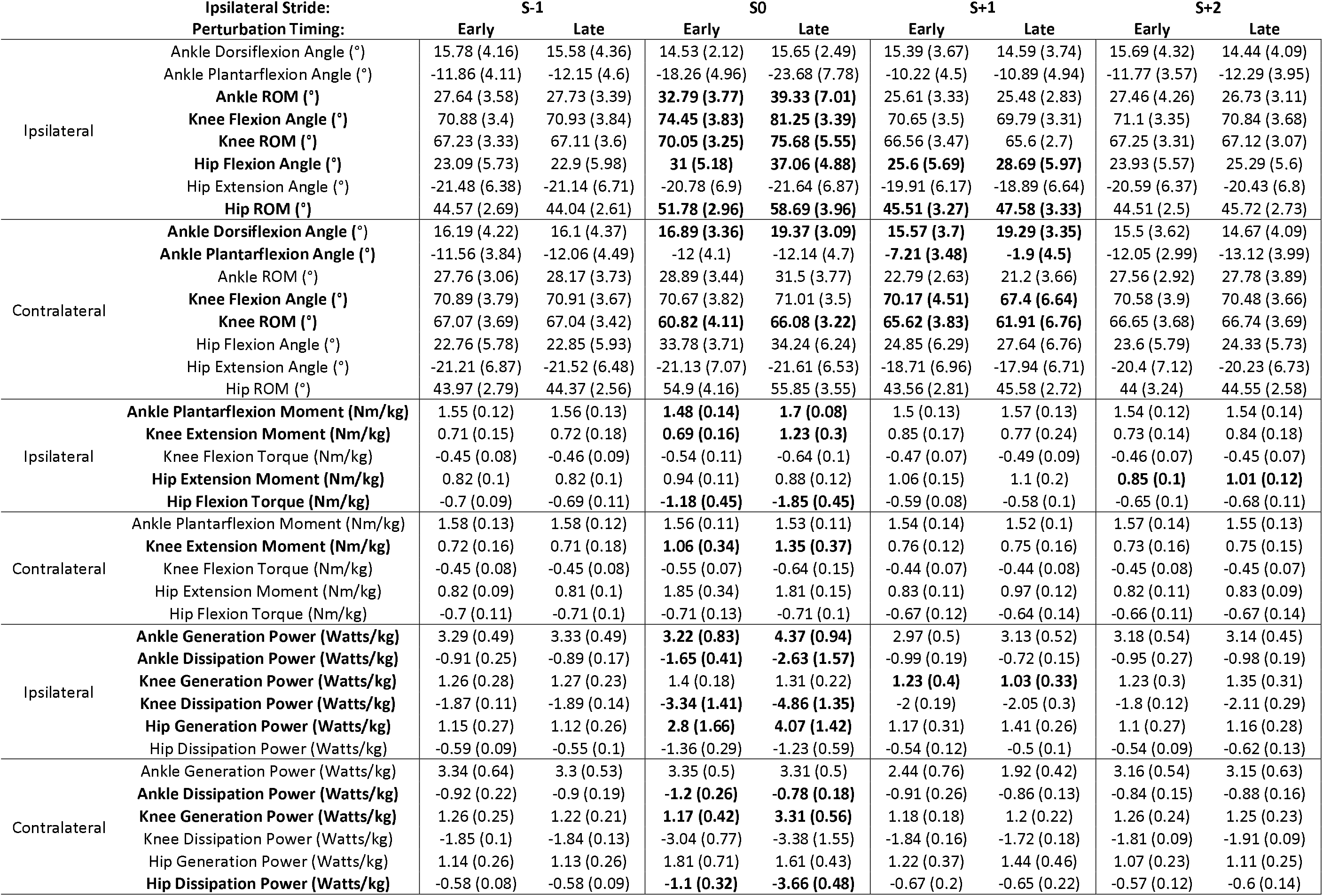
Across-subject mean (standard deviation) peaks in sagittal joint kinematics, kinetics, and mechanical powers. Bold variable names indicate significant timing × stride interactions. Significant post-hoc tests for the effect of timing are indicated by bold values within stride columns.

**Figure S1.**
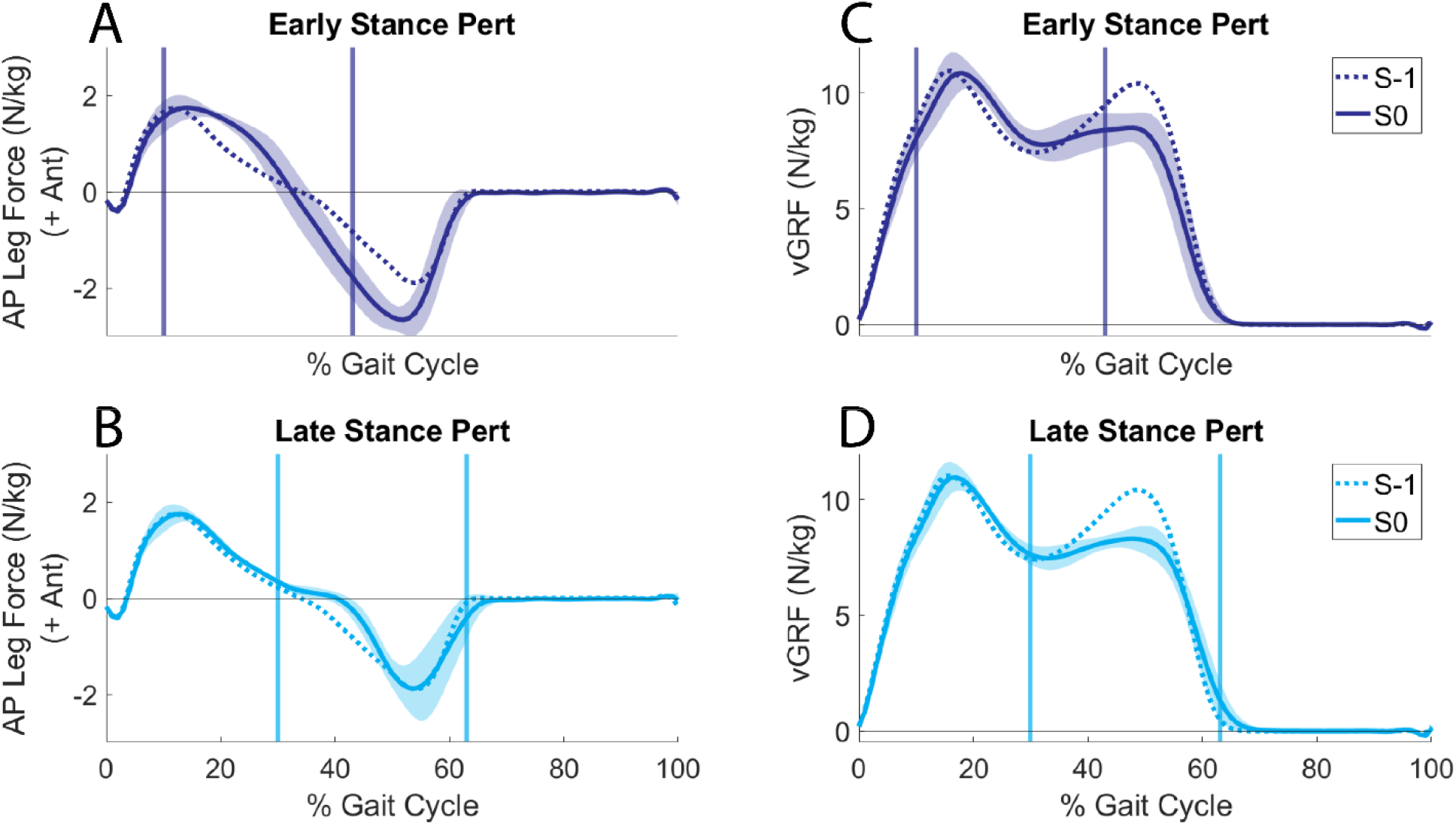
(A-B) Anteroposterior forces exerted by the ipsilateral leg, and (C-D) vertical ground reaction forces of the ipsilateral leg exerted on the COM on the perturbed stride (S0) and preceding unperturbed stride (S-1). Solid vertical lines indicate the average start and end times of the perturbations.

**Figure S2.**
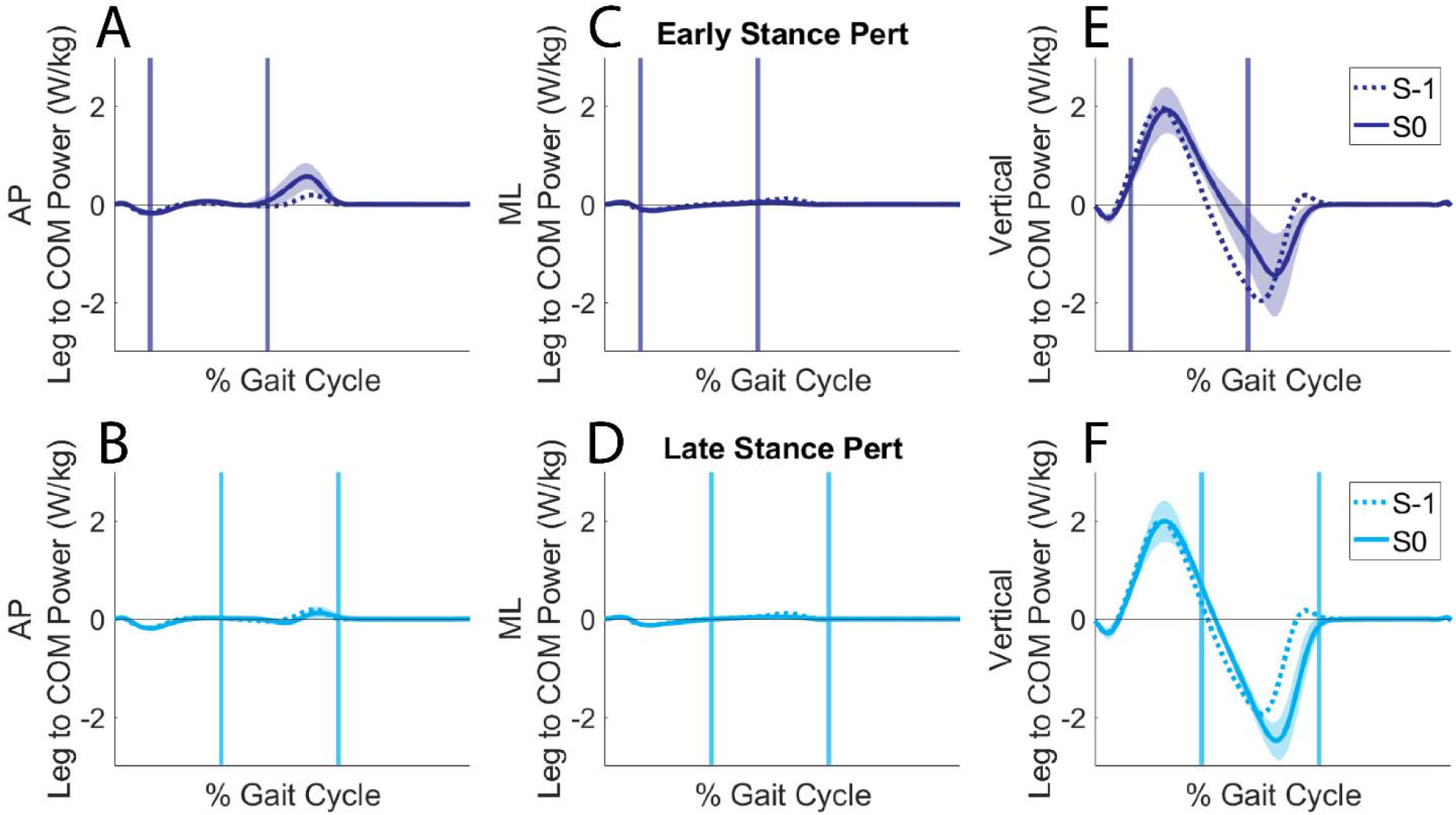
Mechanical power flowing from the perturbed leg to the COM separated into (A-B) anteroposterior, (C-D), mediolateral, and (E-F) vertical contributions during the perturbed stride (S0) and preceding unperturbed stride (S-1). Solid vertical lines indicate the average start and end times of the perturbations.

